# Dynamical Features of Human EEG Scale Systematically with Life Context

**DOI:** 10.1101/125906

**Authors:** Dhanya Parameshwaran, Tara C. Thiagarajan

**Author notes:** Corresponding Author: Tara C. Thiagarajan.

## Abstract

We have previously shown that the complexity of human brain activity and the emergence and strength of alpha oscillation as measured by EEG scale dramatically with access to the features of modernization. Here we show that these features of the EEG, while correlated to one another are distinct in their origin and that while complexity is most strongly related to travel or geofootprint, energy of the alpha oscillation scales more systematically with fuel consumption. Finally, composite EEG scores representing the dominant principal component scale in remarkable lockstep with the dominant component of the features of modernization. This demonstrates that human brain dynamics, and therefore cognitive outcome, are profoundly and systematically tied to the context of life experience. Indeed the implication is that ‘normal’ brain function is not absolute and can only be considered in relation to or conditional on the context in which it is embedded.

## INTRODUCTION

The core of modernization involves technologies and social structures that extend and expand the rate and scope of human interaction and experience. These include agglomeration into dense urban structures, expanded formal education systems and significant technologies such as electricity, telecommunications and motorized transport. Modernization has also brought with it a growing income inequality that results in substantially different levels of access to modern experience. Given that the human brain is an experience dependent organ that continually reconfigures itself in response to stimulus [1-4], such wide differences in experience along these dimensions have a dramatic impact on fundamental aspects of brain dynamics. A host of studies have shown that lower income and education are associated with structural differences in the brain [5-11] and that environmental enrichment in animal studies is associated with a vast array of cellular and molecular changes [12-17]. Further, we have shown in two companion papers that populations with access to higher levels of modernity have significantly higher complexity of the EEG signal [18] as well as dramatically stronger presence of alpha oscillations [19]. Here we present a detailed analysis of the relationship between the features of the alpha oscillation and complexity, and in turn, their relationship individually and in composite with various individual factors of modern context as well as their principal components.

Our sample comprised 402 adults between the ages of 21 and 65 from 48 locations in India including remote settlements of just 300 people with no electricity or motorized transport, to cities of several million people with all modern amenities, and spanned a range of annual incomes from $300 to approximately $150,000 dollars, translating to daily incomes of $0.82 to ∼$410. Correspondingly our sample spanned a spectrum from people living in premodern conditions with no formal education, phones, motorized vehicle or electricity to others who were college educated, digitally connected and internationally traveled. Our measures include the population of the place of residence, income, education, the farthest distance traveled from home (travel or geofootprint) and monthly expenditure on telecommunication, vehicle fuel and electricity (see Table 1 in [18]). Each of these factors represent structures or tools that expand access to interactions and experiences and therefore are proxies for the rate and breadth of exposure of the brain to stimuli.

Here we explore the relationships of each of the EEG features to one another and to individual and composites of each of the contextual features that describe the various components of modern civilization. Our results show that each EEG feature is distinct and has highly significant relationships with many of the contextual features with differing patterns. Finally we show that composite scores of EEG activity derived from principal components of the set of EEG features move in remarkable lockstep with composite scores of modern experience derived from principal components of the set of contextual features demonstrating a very clear relationship between brain dynamics and life context.

## MATERIALS AND METHODS

### Sampling, Survey and EEG Recording

The survey methodology and sampling were carried out as described in a companion paper [18]. Briefly, participants answered a series of questions regarding their demographic, communication and mobility behavior in addition to having EEG recorded for three minutes when they were seated with their eyes closed. All participants were explained the purpose and protocol of the experiment and signed an informed consent. The EEG recordings were carried out using the Emotiv EPOC device, after comparison to the clinical grade Neuroscan device for similarity of results, again as described in [18].

### Principal Component Analysis

Principal Component Analysis was done using the FactomineR PCA function. Prior to application of the PCA function, all records without values in any one of the columns were removed from the analysis (121/402 records). Prior to performing the PCA all data components with highly skewed distributions approximating lognormal were log transformed with log base 5 to create linear relationships among the variables. This included Income, Fuel Spend, Phone Spend, Electricity Spend and Population among the contextual variables, and E_α_ from the EEG features. The circle of correlations shown represent the unit scaled coordinates of the components. The component scores for the EEG and context of each individual were calculated as the sum of the contribution of each element multiplied by the unscaled individual element.

### Statistical Significance of Trends

All trends are shown as population means ±SEM for each bin along the ordinate axis thereby depicting a trend of the shift in the population distribution. We calculated various statistics to determine the significance of these trends. First we calculated the R^2^ of logarithmic, exponential and linear fits, reporting the fit with the highest R^2^ value. We next computed the significance of an ANOVA (*p_ANOVA_*), which would provide the probability of a difference across the various bins. To determine the likelihood of such a trend appearing that could not be accounted for by intraperson fluctuations, we also computed the probability of finding a similar trend from shuffling the C_T_ values across the participants 1000 times including: (i) the probability of obtaining a significant ANOVA in the shuffled iterations that was ≤ 0.05 (*p_skuff1_*) or (ii) ≤ the *p*-value of the data (*p_skuff2_*) and (iii) the probability of finding a shuffled iteration with a trend that was positively correlated with *p*<0.05 to the data (*p_skuff3_*).

## RESULTS

### Relationships among EEG Features

Here we look at the relationship among the three EEG features to determine how they are related. It is possible that the various features of the EEG signal are different aspects of a common underlying influencer and therefore strongly correlated. On the other hand given that an oscillation is a repetitive structure and complexity is a measure of diversity, a strong alpha oscillation (i.e. with high energy E_α_ and high peak frequency P_α_) would reduce the complexity of the structure and may therefore be negatively correlated with complexity.

Fig. 1A shows a scatter plot of C_T_ vs. P_α_ for all participants in our study along with the mean C_T_ ±SEM (in black) for the population in 0.5 Hz bins of P_α_. Contrary to the expectation, mean CT increased systematically with P_α_ (R^2^ of a linear fit of the population means was 0.61 with slope = 1.78). However the overall correlation between P_α_ and C_T_ was positive but very low (r =0.2). Indeed individuals had a broad spread of CT values for each value of P_α_ and a participant with no detectable oscillation could have a CT value spanning the entire range of possibilities from 35 to 90. The structure of this relationship indicates that these two features are likely to be independent in their origin though may be influenced by some common drivers.

**Figure 1.**
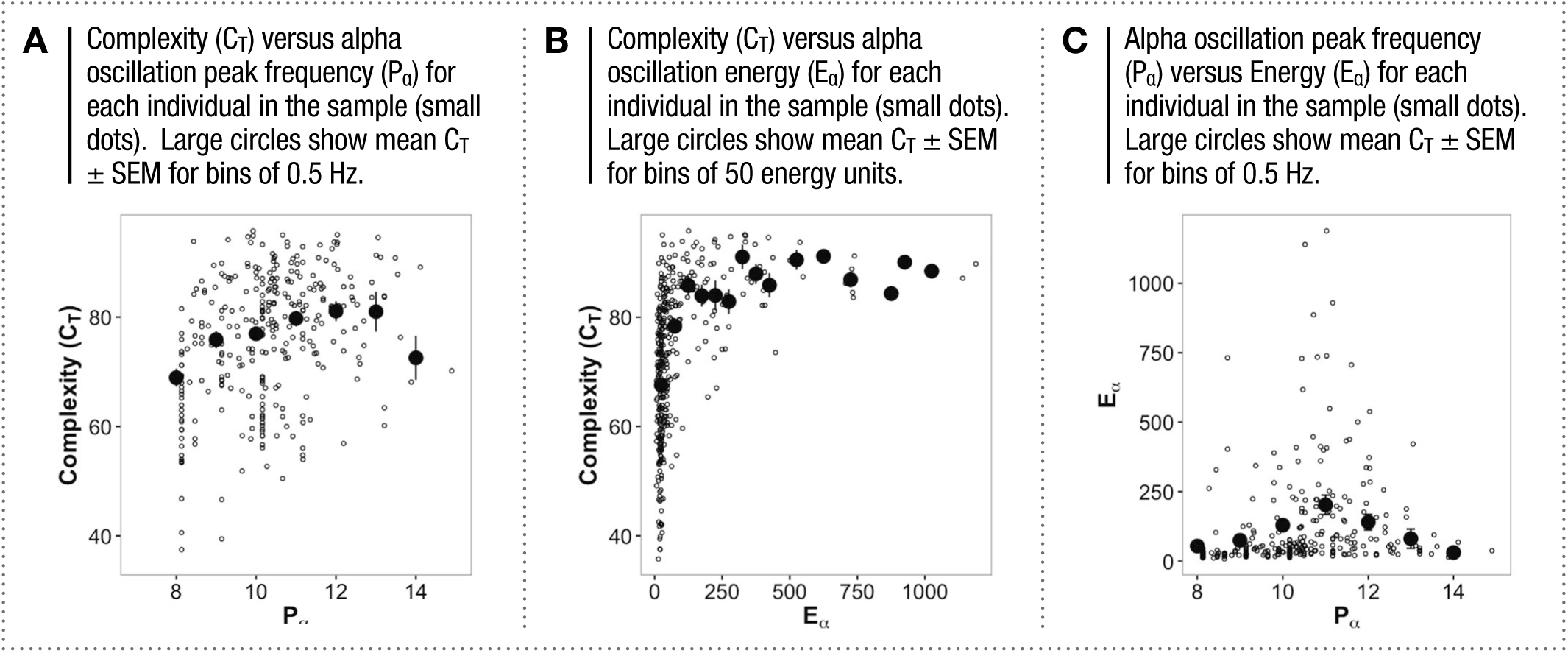
RELATIONSHIP BETWEEN EEG FEATURES

We next looked at the relationship between C_T_ and E_α_ (Fig. 1B). Here again the correlation of the two features across all participants was positive but weak (r = 0.43). People with very low values of E_α_ reflecting the absence of an oscillation, had C_T_ values spanning the entire range. However unlike the case of P_α_, while the range of CT values was higher for those who did have the oscillation (and therefore E_α_ values above 40), the population means did not change substantially as E_α_ increased beyond this level (R^2^ of linear fit = 0.33, Slope = 0.01) indicating, again, distinct origins and influencers of complexity and the features of the alpha oscillation.

Finally we note that the two features of the alpha oscillation, P_α_ and E_α_ are also only very weakly correlated (r= 0.09, Fig. 1C) and therefore represent two distinct features of the oscillation. Indeed while P_α_ is distributed normally, E_α_ has a substantially different long tailed distribution demonstrating distinct structures and origins. This relationship is shown in our previous paper (Fig. 3E; Alpha Energy). Mean E_α_ had a U-shaped relationship to P_α_ with the highest values at ∼11 Hz in the middle of the alpha range. This U-shape could not be accounted for by a boundary effect of the range suggesting that there may be an optimal P_α_.

**Figure 3.**
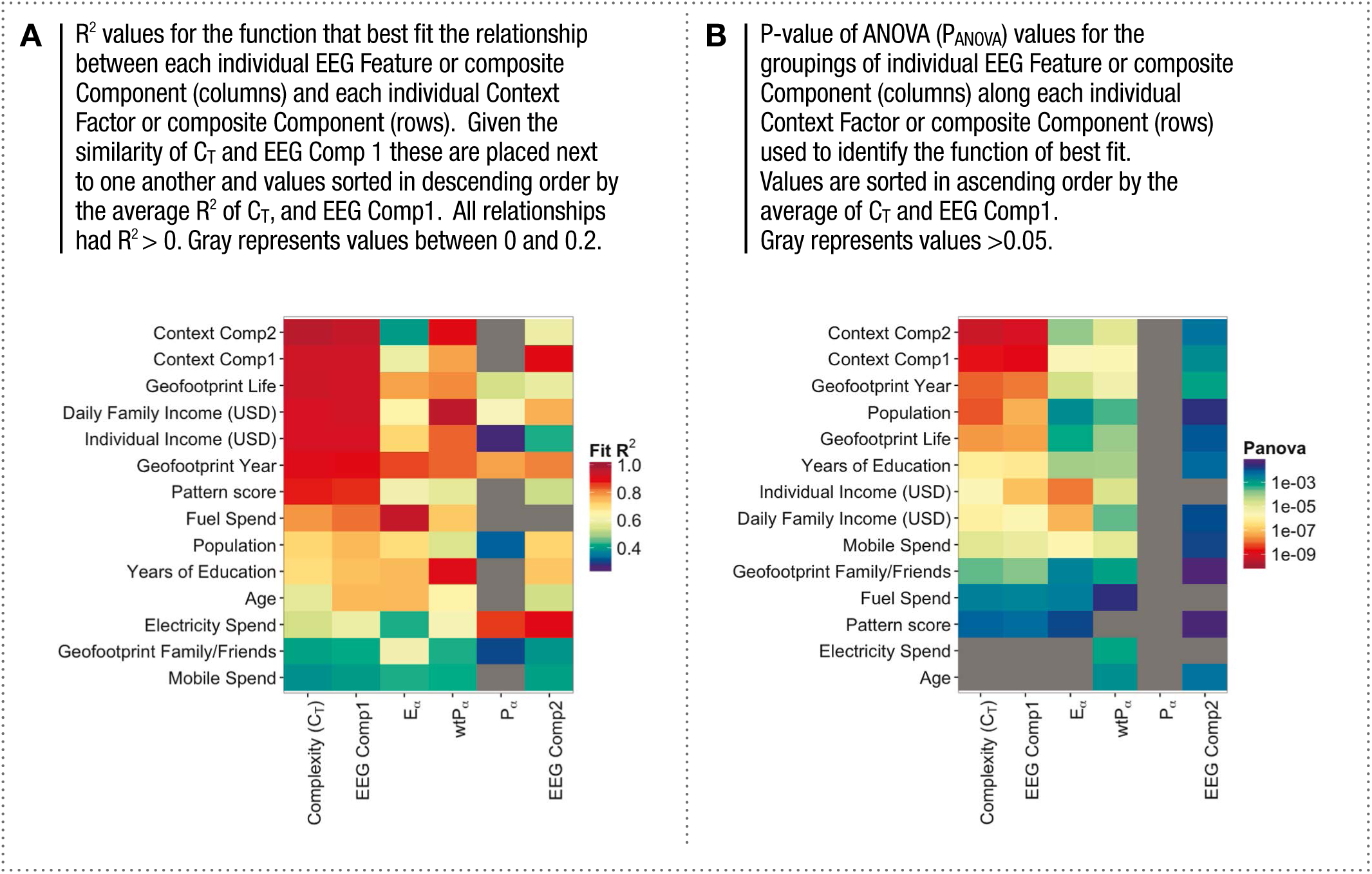
SUMMARY OF RELATIONSHIPS BETWEEN EEG FEATURES AND CONTEXT FACTORS

### Principal Component Analysis of EEG Features and Context Factors

We next performed principal component analysis to identify orthogonal components among both the EEG features (Fig. 2A,B) and the contextual factors (Fig. 2C,D) of the participants. Contextual factors refer to the life factors such as education, technology use and others that we have captured; essentially the context in which the participant lives.

**Figure 2.**
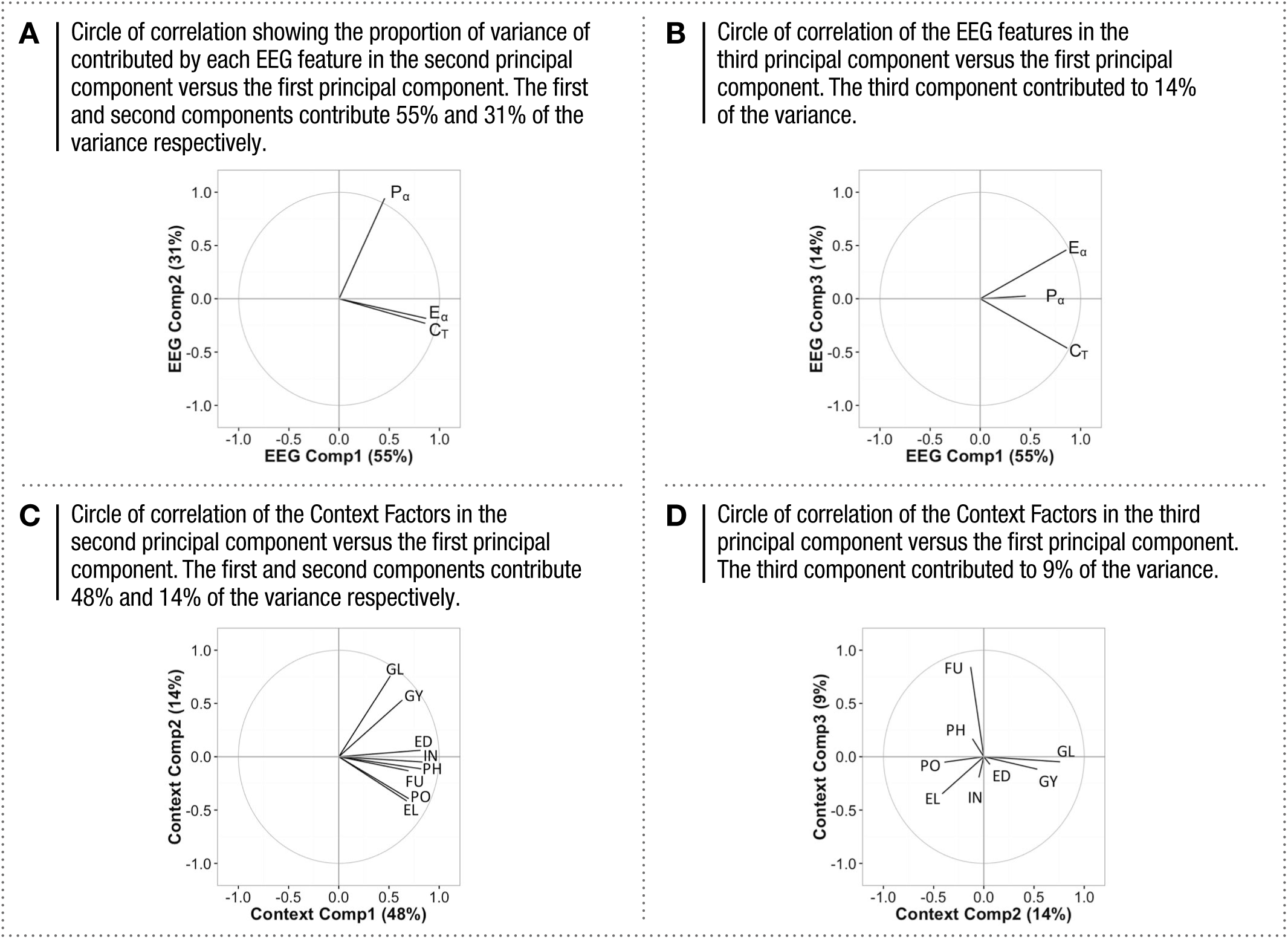
PRINCIPAL COMPONENTS OF EEG FEATURES AND CONTEXT FACTORS

The first component of the EEG features explained ∼55% of the variance in the sample and represents the correlation of all three features. The second component explained ∼30% of the variance and the third component the remaining 14%. The contribution of the factors within the first principal component relative to the second is shown as a vector (circle of correlations) with one component represented on each of the x- and y-axes. Fig. 2A shows the second component of the EEG features (EEG Comp 2) versus the first (EEG Comp 1), and 2B shows the third component (EEG Comp 3) versus the first. These demonstrate that the second component largely represents the residual dominance of P_α_ and the third component the residual variance of E_α_.

In the case of contextual factors, the PCA was performed using 8 factors. While all factors were positively correlated, we have previously shown that income, education and phone spend were most tightly correlated (Fig. 4A in [18]) while travel or geofootprint was least correlated to the other factors. Correspondingly, the first principal component (Context Comp 1) accounted for 47.54% of the variance where all factors of experience were essentially positively correlated with income, while the second component (Context Comp 2) contributed 14.24% of the variance and was dominated by travel (Geofootprint) (Fig. 2C). The third component (Context Comp 3), shown here versus the second component, was dominated by the residual variance of expenditure on vehicle fuel and accounted for only ∼9% of the variance (Fig. 2D). We note that fuel expenditure is a proxy for more local activity and also the speed with which the environment might be experienced, while geofootprint represents exploration of new environments including navigating new spatial layouts, languages and cultures.

**Figure 4.**
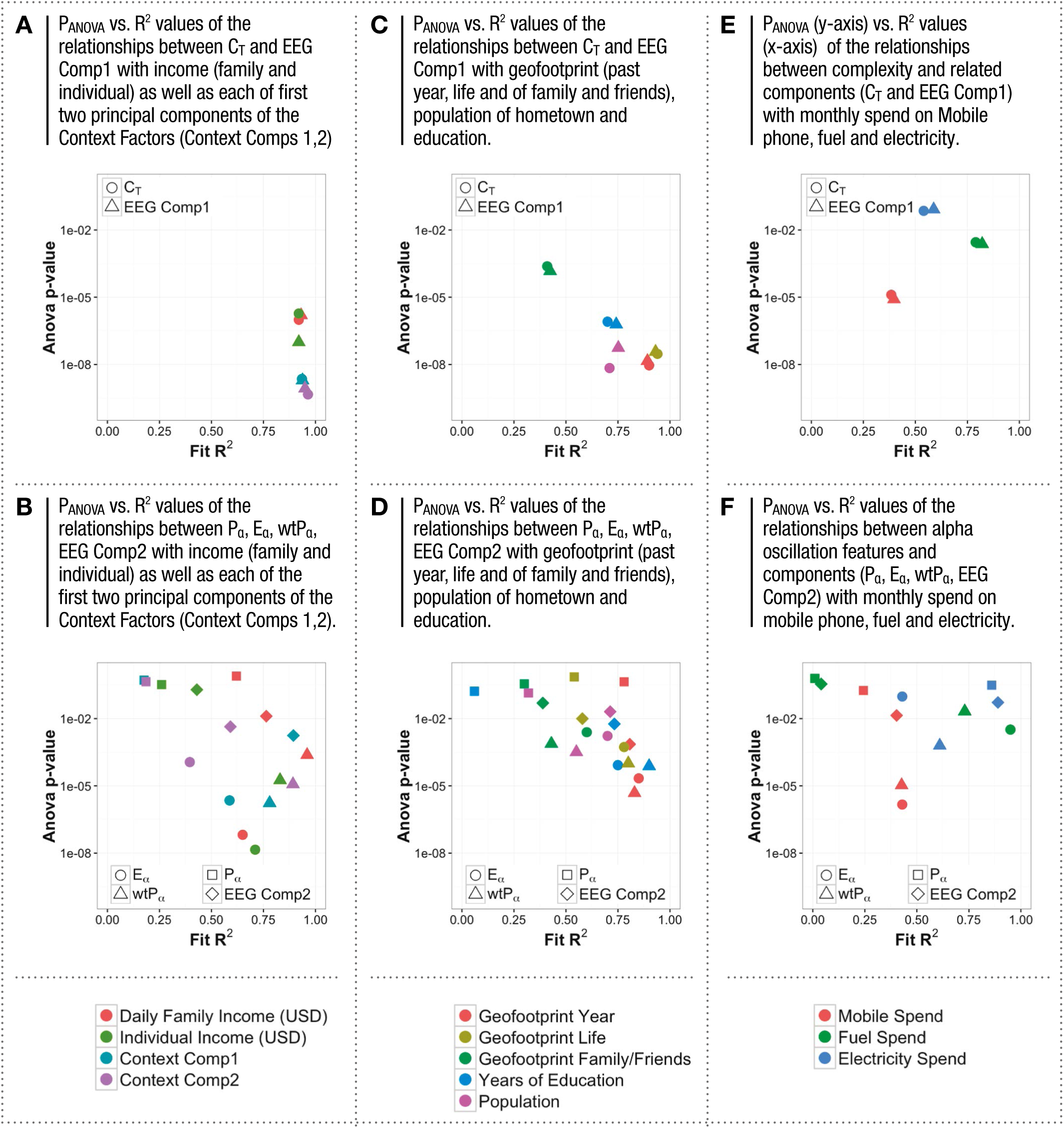
R^2^ vs. *P_ANOVA_* FOR EEG FEATURES AND CONTEXT FACTOR

Going forward, along with the individual EEG features we describe the relationships of the scores for each of the first two principal components of the EEG features to the scores for each of first two components of the contextual factors.

## RELATIONSHIPS BETWEEN EEG FEATURES AND CONTEXT FACTORS

Here we compared 6 aspects of the EEG signal (EEG features C_T_, P_α_, E_α_, weighted P_α_ and composite principal component scores EEG Comp1 and Comp2) to 14 contextual or behavioral factors and components (11 features of modernization, Pattern Test Score and the first two principal components of the context factors Comps 1 and 2). Given that this results in 6x14 or 84 comparisons, we present in the main paper only an overview of the strength and significance of all the relationships (R^2^, significance of an ANOVA *P_ANOVA_*; Figs. 3,4) as well as some of the most significant and interesting relationships (Fig.5,6). However, details of all the relationships for each of 8 context factors are shown in 9 supplementary figures.

**Figure 5.**
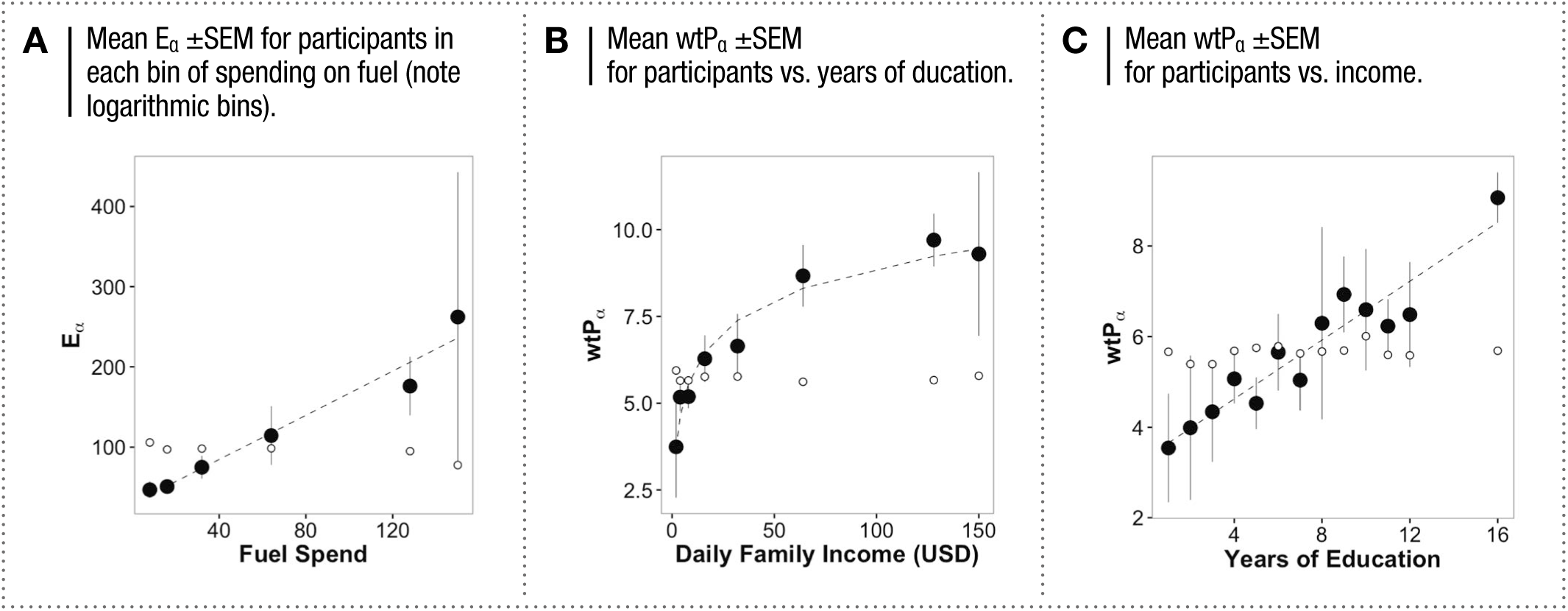
KEY RELATIONSHIPS OF ALPHA OSCILLATION FEATURES TO CONTEXT FACTORS

**Figure 6.**
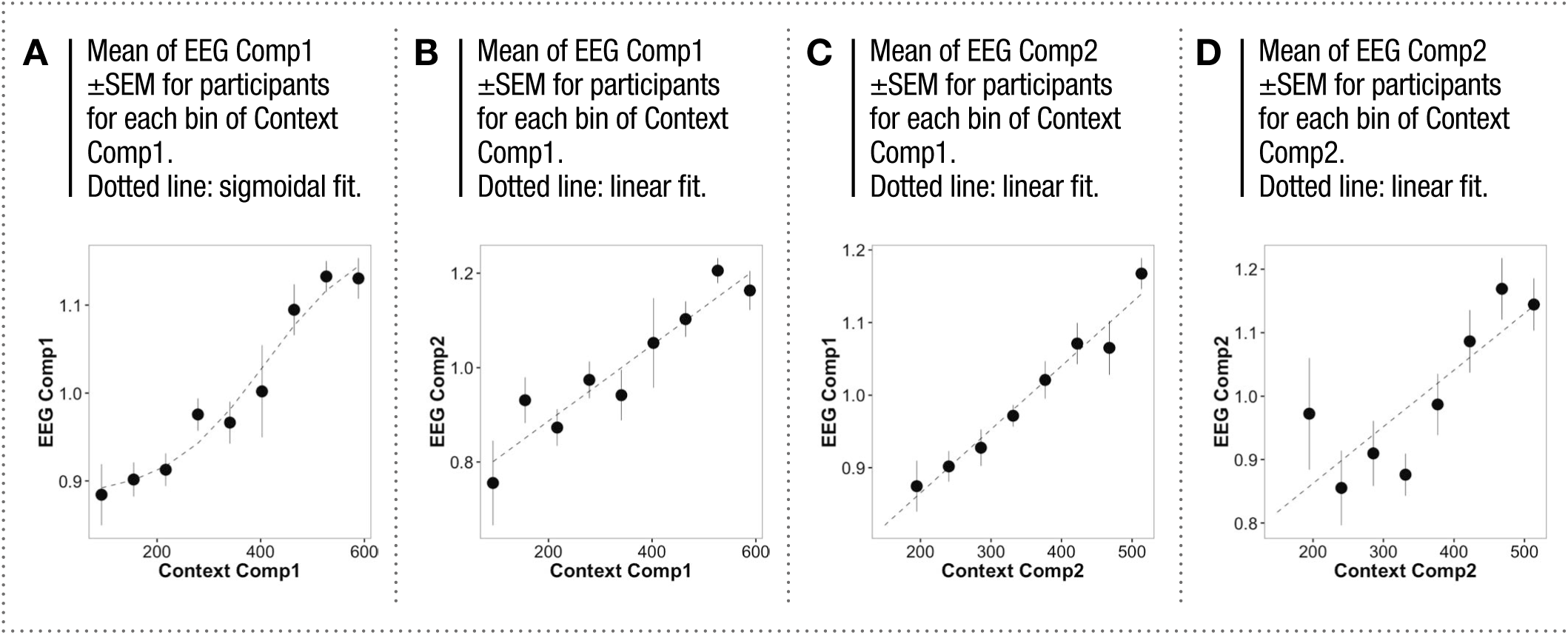
SCALING OF COMPOSITE EEG FEATURES WITH COMPOSITE CONTEXT FACTORS

To compute the R^2^ and *P_ANOVA_* we performed an analysis of the trend with which the population means of the EEG feature or component score shifts as a function of an individual context factor or component score. The bin sizes for the context factors were determined such that the first and last bins (with lowest and highest value), where the number of participants was typically fewer, had at least 5 data points. We then fit trend lines through the means and calculated the R^2^ for the function with the best fit. Typically the functions of best fit were either linear or logarithmic, although some exponential and sigmoidal relationships were found. We then performed an ANOVA for the groupings, calculating the p-value (*P_ANOVA_*). The R^2^ and *P_ANOVA_* together provide views of the magnitude and significance of the trend. We also computed various probabilities of positive trends and *P_ANOVA_* values for shuffled datasets (see methods). All statistics for the 25 most significant relationships are shown in Table 1. The comprehensive statistics for all 84 comparisons are in Supplementary Table 1.

**Table 2.**
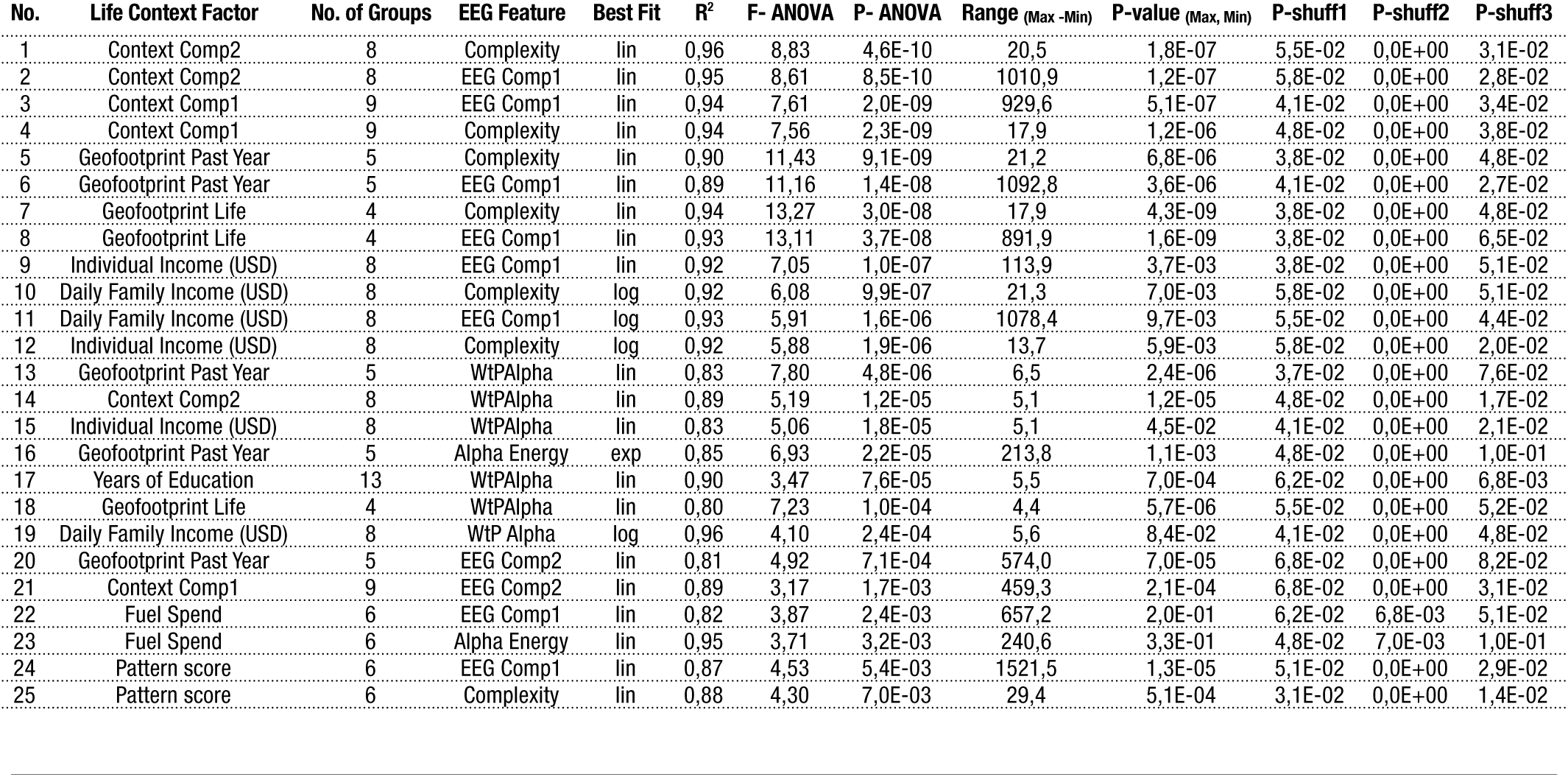
STATISTICS FOR MOST SIGNIFICANT RELATIONSHIPS

Fig. 3A shows the R^2^ of the best fit for each context factor versus each EEG feature. Along the x-axis the matrix is sorted by the correlation of the EEG features/principal components to each other thereby bringing C_T_ and EEG Comp1 next to one another. Along the y-axis the EEG features are sorted based on the average R^2^ of CT and EEG Comp1, which tended to be similar as CT dominated EEG Comp1. The strongest and most significant relationships, with R^2^ >0.9, were between each of C_T_ and EEG Comp1 and the composite component scores of the factors (Context Comp1, which was roughly evenly made up of all factors and Context Comp2, which was dominated by Geofootprint). The next strongest were between each of C_T_ and EEG Comp1 and Geofootprint and then Income, which had R^2^ values above 0.8. We note here that given that geofootprint represents an ordered categorization of an underlying variable, but not a systematic or ordinal scale, a fit should strictly not be performed. However we have included it for the purpose of approximate comparison.

With regard to the features of the alpha oscillation, the largest R^2^ was between E_α_ and fuel consumption or fuel spend (0.95, exponential fit). In contrast the fits for P_α_ relative to the context factors were not as strong, although wtP_α_ (P_α_ weighted by E_α_) had strong relationships with income as well as the composite Context Components scores, particularly Context Comp2.

Fig. 3B shows the *P_ANOVA_* for the grouping of EEG features into each bin of the context factor of comparison (as defined in 3A above). Here again, the matrix is sorted along the x-axis in the same way as in 3A. The y-axis is sorted by the average *P_ANOVA_* of the first two columns (C_T_ and EEG Comp1) going from the lowest average *P_ANOVA_* (most significant) to the highest (least significant). Note that the ordering is therefore not the same as in 3A and that grey indicates a value of 0.05 or greater. The most significant relationships also had the best R^2^ values for trend fits. Prominently, the relationship between EEG Principal Components and Context Principal Components had PANOVA values of 10^−9^ or lower indicating a profoundly strong relationship between dynamical features of the brain and life context.

In order to see both components of the relationship R^2^ and *P_ANOVA_* together we plotted these against each other (Fig.4). Figs. 4A, B shows the R^2^ versus *P_ANOVA_* for each of C_T_ and EEG Comp1 (4A) and the Alpha Oscillation features (4B) to Context Comp1, Context Comp2, Daily Family Income and Daily Individual Income. Both income and the component scores are more macro factors. While the component scores are a weighted score of all the factors, income is an enabler of all the factors.

Figs. 4C,D show the similar relationships for Geofootprint, Population and Education which are demographic and environmental factors of exposure. In this group Geofootprint had the greatest R^2^ and significance with respect to most EEG features, more so than education. The most significant relationship was with complexity and the first component score as shown by their position in the bottom right corner with R^2^ >0.85 and pANOVA <10^−7^.

Fig. 4E,F shows the relationship of EEG features with expenditure on technology; mobile phone, fuel and electricity. Relative to the other categories, technology factors had far less significant relationships to these EEG features and only mobile and fuel spend fell below the significance level of 0.05 (dotted line). Interestingly, E_α_ had a more systematic and significant relationships to mobile phone and fuel spend (Fig. 5B,C) than the C_T_ and EEG Comps 1 and 2. The relationship between fuel spend and E_Œ_ was the most significant stand out. The R^2^ was very high though the *P_ANOVA_* was only moderately significant (P=0.003). This was on account of a systematic skewing of the distributions rather than the progressive shift that was characteristic of geofootprint and education.

In Fig.5 we show the most noteworthy relationships between the EEG alpha oscillation features and individual factors: between E_α_ and fuel consumption (5A), and wtP_α_ and education (5B) and income (5C). Note that the relationships of Complexity with these factors has been shown in [18] and has therefore not been repeated here. The complete set of relationships between all EEG features and Context factors are shown in Supplementary Figures: Daily Family Income (Sup. Fig. 1), Education (Sup. Fig. 2), Geofootprint/Travel (Sup. Fig. 3), Mobile Phone Usage (Sup. Fig. 4), Fuel Consumption (Sup. Fig. 5) and Electricity Spend (Sup. Fig 6).

Finally, we show the relationships between the first two EEG Components and the first two Context Components (Fig. 6). As described earlier, the EEG components represent composite views of the overall dynamical characteristics along the components of maximal variability among the population. The first EEG component (EEG Comp1) scaled sigmoidally in relation to the first Context Component (Fig. 6A; *P_ANOVA_*= 1.96 × 10^−9^ R^2^=0.94) and linearly with the second Context Component (Fig. 6B; *P_ANOVA_*= 8.54 × 10^−10^ R^2^=0.95). The second EEG Component (dominated by P_α_) was not as tightly related but nonetheless scaled linearly with both Context Components 1 and 2 (Fig. 6C,D; *P_ANOVA_* = 1.75 × 10^−3^ R^2^= 0.89 for Context Comp1 and *P_ANOVA_*= 4.54 × 10^−3^ R^2^ = 0.59 for Context Comp2). However, we do point out that this does not imply a linear relationship with the components of modern life. Given the long tail structure of the various data elements such as income and mobile phones, the principal components are derived from logarithmic transforms of many of the data elements. Thus the effective relationships are nonlinear in nature.

## DISCUSSION

We have described here the relationship between three fundamental aspects of the resting EEG signal, the complexity of the waveform and the peak frequency and energy of the alpha oscillation, and the contextual factors of income, geofootprint, population, education and spending on mobile phone usage, fuel and electricity. Overall we show that the magnitude of these EEG features increases as a function of these life context factors indicating that the more access one has to the various aspects of modern living, the more enhanced these features were in the brain.

A host of studies have shown that lower income and education are associated with structural differences in the brain [5-11] and that environmental enrichment is associated with a vast array of cellular and molecular changes in animals [12-15]. However this study is the first to combine observations of EEG with such a large number of contextual factors and such a broad crosssection of human populations. Our results provide not just a deeper understanding of the various factors of influence and their interrelationships, but also a clear picture of the magnitude of difference and the manner in which the change occurs. Many of the relationships were highly systematic and of unquestionable magnitude and significance. Overall, complexity and the first principal component EEG Comp1, which was dominated by complexity, had the most dramatic relationship to the various contextual factors of modernization. We discuss below some of the key findings and their implications.

### Geofootprint as a key influencer

Every EEG feature except P_α_ had a dramatic and highly significant relationship to geofootprint or the spatial extent of one’s travel, with the relationship with Complexity being the most systematic. Further, geofootprint reflecting travel in the past year appeared to have higher significance than the geofootprint of ones lifetime. This ran counter to our expectation that education would have the most dominant correlation to ones brain complexity. In this context it is interesting to note that while education and mobile phone use were very tightly coupled to income, geofootprint varied more independently of income and other factors. This makes sense when one considers that unlike education, which is an institutionalized experience, geofootprint or the exploration of the physical world generally comes about by personal choice. Such exploration is also a complex interactive experience compared to the more didactic, text book based learning that is characteristic of the Indian education system. Traveling requires planning and navigating new spatial environments, people, languages and cultures and a forced learning of new things. Indeed, migratory populations that have traveled long distances in search of new resources have played a fundamental role in shaping human history. It will be of considerable interest to explore more deeply the nature and direction of influence of travel and migration.

### The impact of education

In our study, years of education like geofootprint, was also significantly related to all EEG factors except P_α_, although with slightly less significant statistics. Overall, we see that the enhancement of EEG features is most prominent for the first five years of education while the effects of the years of secondary education are not very significant. However, those who make the transition to college are considerably higher in their EEG metrics perhaps because they are a select group that has performed at a higher level in their secondary education. This group also has significantly higher income overall, as they could not otherwise afford to go to college. This also means that their secondary education is likely to have been of higher quality. Finally, many of our participants were from small villages and towns in India where the method of education was purely textbook based without the interactive technology and tools of today. It would therefore be of interest to explore different methods of education to determine how much our results are a consequence of a poor quality of education or a particular system rather than a general statement about classroom instruction.

### The logarithmic effect of income

Both complexity and the weighted P_α_ had very strong relationships with income, which is one of the most significant enablers of access to experiences and technologies. In both cases the relationship with income was clearly logarithmic. The steepest part of the curve was between $1 and $30/day in family income indicating that there were substantial gains as income increased up to $30/ day. However as one went beyond $50/day the differences in the populations were no longer substantially different. $30/day corresponds to an annual income of $10,950, which in India provides access to all features of modern living. In PPP terms $30/day in India is roughly $90/day in the United States and is considered high income by global standards. At this income level, one can comfortably own a car, smart phone and modern household amenities as well as have 24 hour internet access. Importantly, in 2011 PPP terms, globally only 7% of humanity live on >$50 a day (considered high income) and 16% on $20-$50/ day (considered upper-middle income) respectively; in India only about 0.5% have incomes between $20 and $50 per day[20]. By contrast in the United States 89% have incomes >$20/day.

It is also of interest to note that a study in the United States showed a similar logarithmic relationship to the surface area of the cortex[9]. Indeed the various structural differences in the brain that are associated with brain structure such as increased cortical thickness and surface area [5-11] would be expected to translate to changes in its aggregate dynamical function as measured by EEG.

### Fuel consumption and the alpha oscillation

Compared to complexity, the features of the alpha oscillation were not as systematically related to any of the features of modernization that we looked at. However, it is clear that the alpha oscillations are much more prominent in the higher income groups and were virtually undetectable in the majority of the premodern group. Of all the relationships, the most intriguing was between the energy of the alpha oscillation and fuel consumption. Unlike geofootprint, which captures a more complex experience of navigating new places, fuel consumption reflects largely daily local car travel. Consequently it is also a proxy for the speed at which a person experiences visual stimuli. It has been suggested that the alpha oscillation plays a role in enabling attention on salient aspects while suppressing others [21, 22]. This would therefore be a useful feature that was substantially enhanced as we began experiencing the world at faster speeds.

### Implications

These metrics of the EEG that we have shown clearly scale with various elements of life context. What does this mean in the context of cognitive function? The alpha oscillation has been widely studied and demonstrated to have positive correlation to a number of cognitive capacities[23, 24] such as working memory[25-27], attention[28-30] and learning ability[31]. Conversely it declines with increasing age[27] and is found at lower levels in fragile X mental retardation syndrome[32] and in patients with Alzheimer’s and amnesic mild cognitive impairment[33]. Complexity, as we have defined it, has not been extensively tested in its relevance to cognitive function. However, our limited study has shown a very significant correlation to performance on a pattern completion test [18]. Various entropy measures, however, which are closely related to this measure of complexity correlate with levels of consciousness measured during the application of anesthesia [34, 35] as well as by comparing people who are in different states of consciousness (e.g. minimally conscious and coma)[36]. In the context of our findings this suggests that modern advances have dramatically enhanced cognitive function. More poignantly our findings indicate that as a species, we are diverging dramatically in our cognitive function. Unlike the heart, which beats and performs in the same way for all of us, our brain does not. Indeed, the metrics of brain dynamics can only be described in context and there is no absolute ‘normal’ like there is for other organs such as the heart. With the tools of modern advance and the its correlates of cognitive function still a relative privilege for much of humanity, what does this mean for our future as a collective species?

## ACKNOWLEDGEMENTS

A very special thank you to Sathish S, Aravind and Govind for managing the field EEG recordings and surveys, as well as S. Ravi Shankar who enabled smooth coordination of the process. We also thank the various staff at Madura Microfinance who facilitated access to the villages and recruitment of participants and SciSphere, India for providing us survey tools, access to demographic and economic data for location selection and data support.

---

**Sup. Figure 1.**
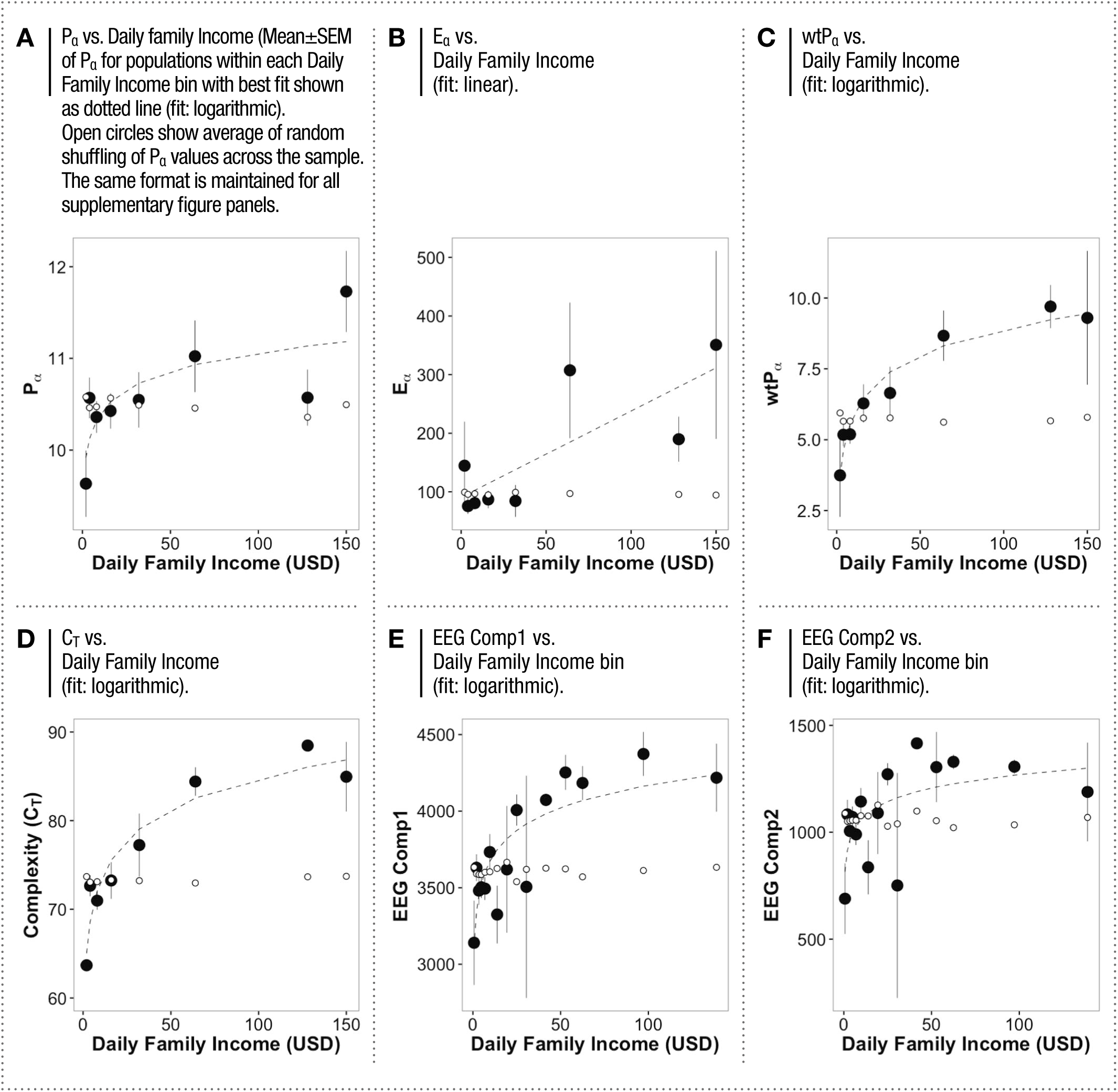
RELATIONSHIPS OF EEG FEATURES TO FAMILY INCOME

**Sup. Figure 2.**
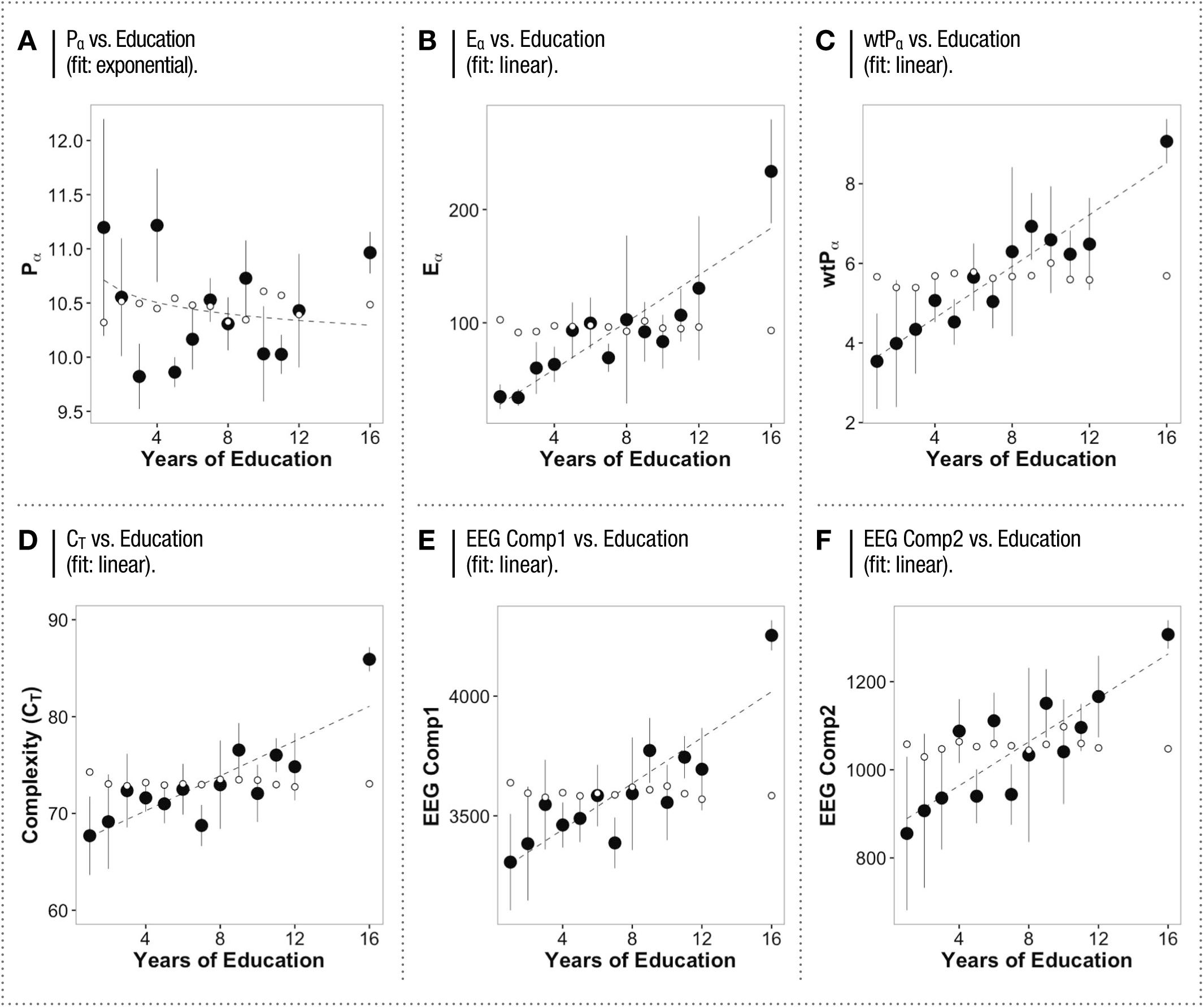
RELATIONSHIPS OF EEG FEATURES TO EDUCATION

**Sup. Figure 3.**
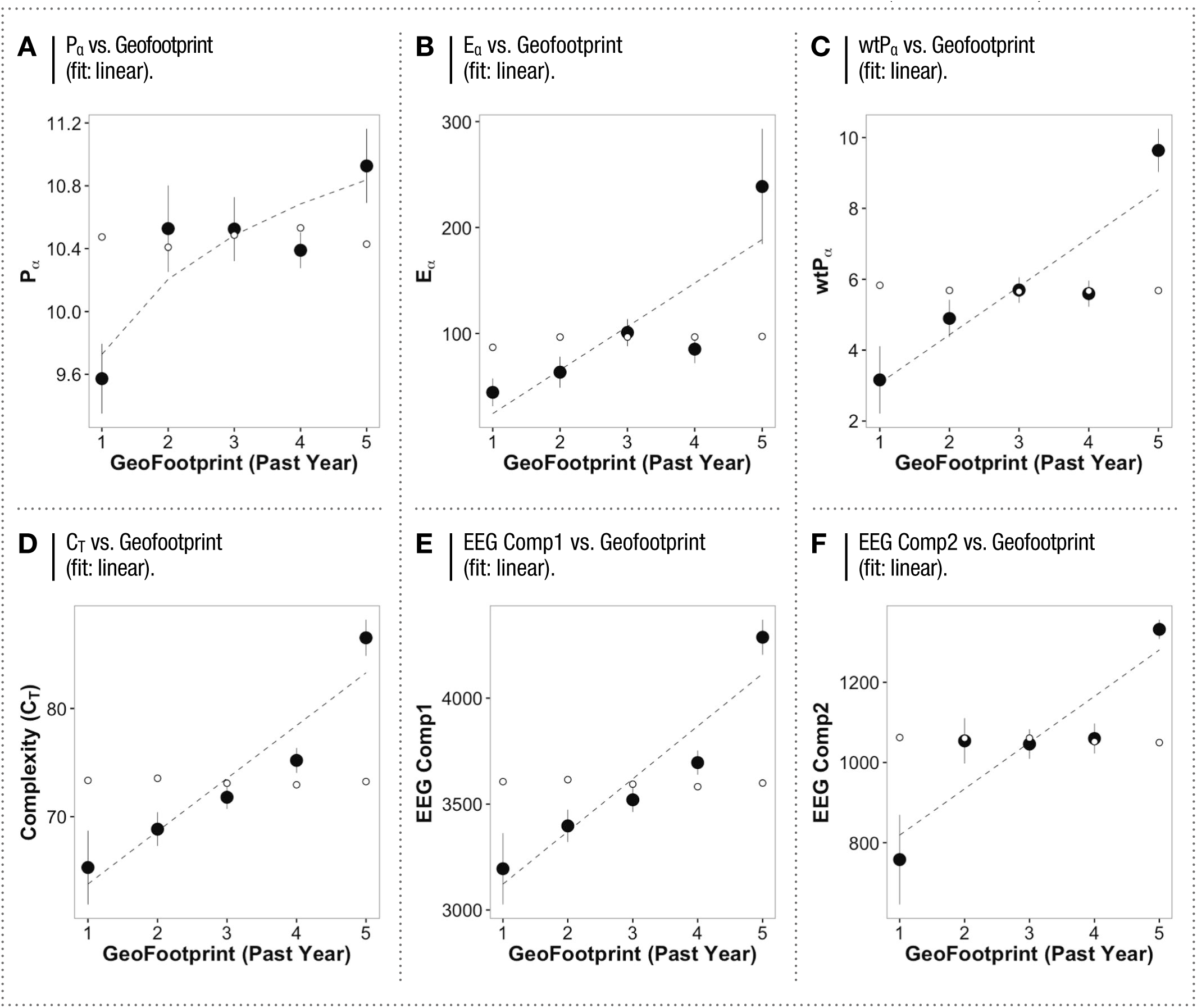
RELATIONSHIPS OF EEG FEATURES TO GEOFOOTPRINT (PAST YEAR)

**Sup. Figure 4.**
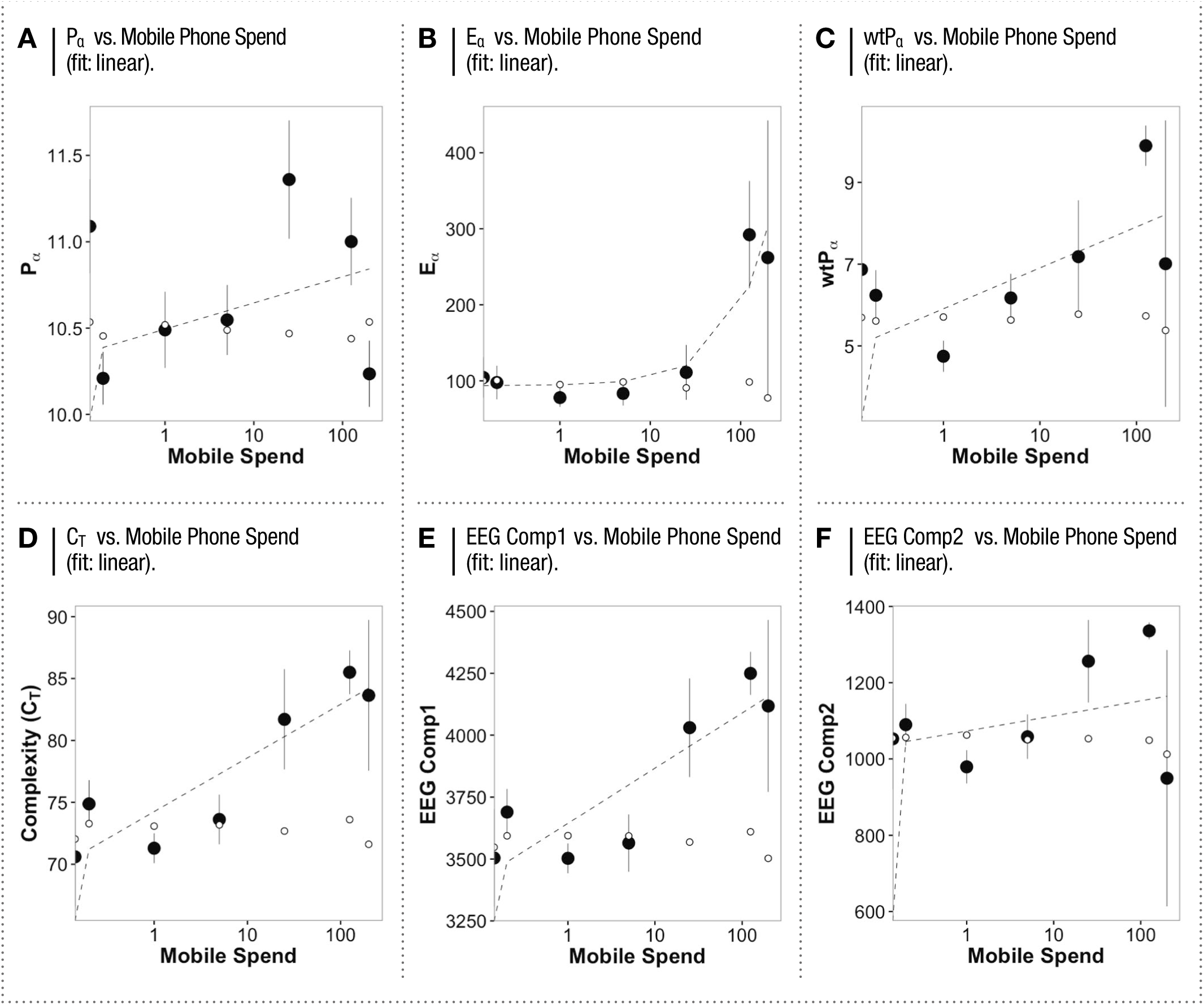
*RELATIONSHIPS OF EEG FEATURES TO MOBILE PHONE SPEND*

**Sup. Figure 5.**
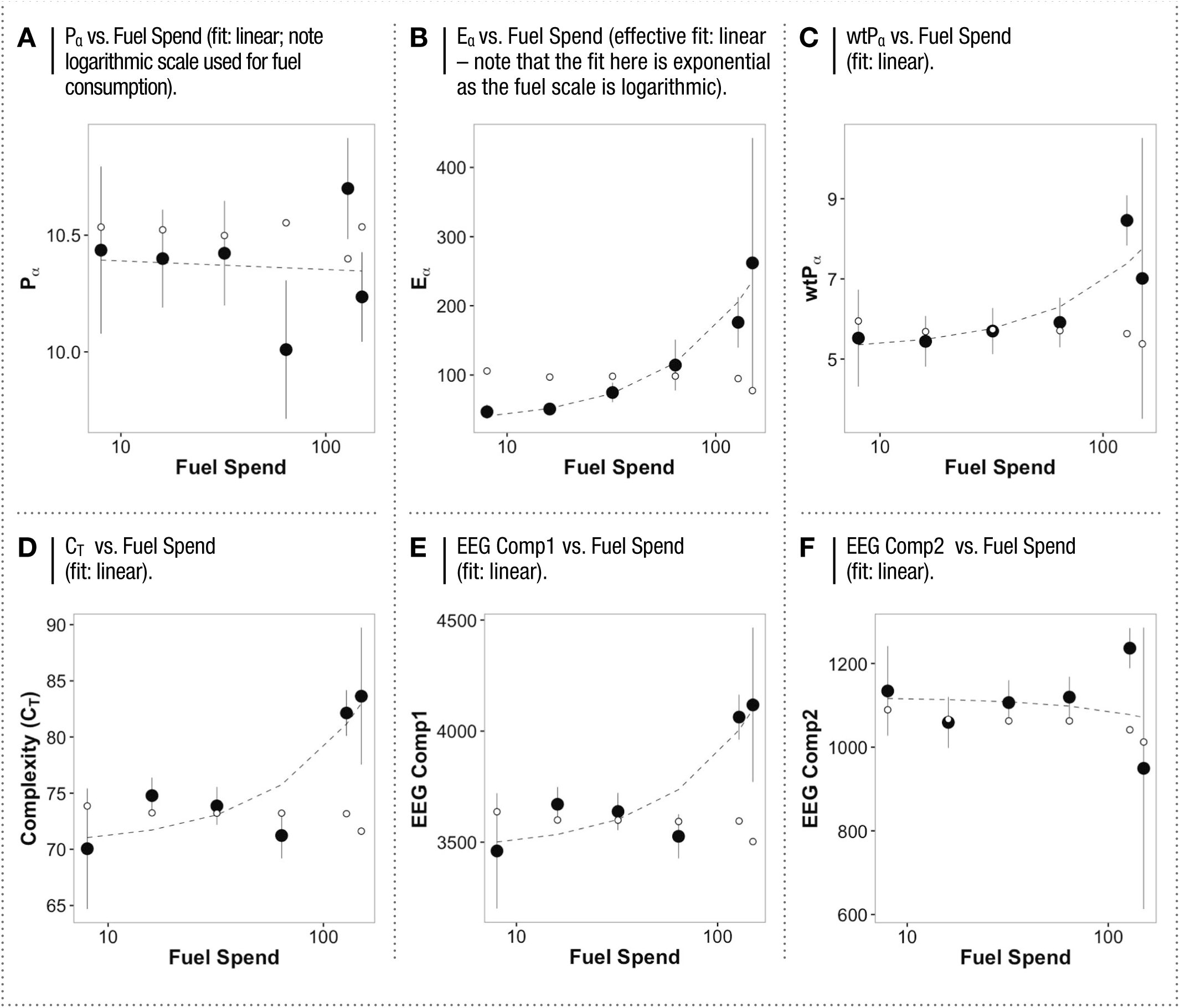
*RELATIONSHIPS OF EEG FEATURES TO FUEL SPEND*

**Sup. Figure 6.**
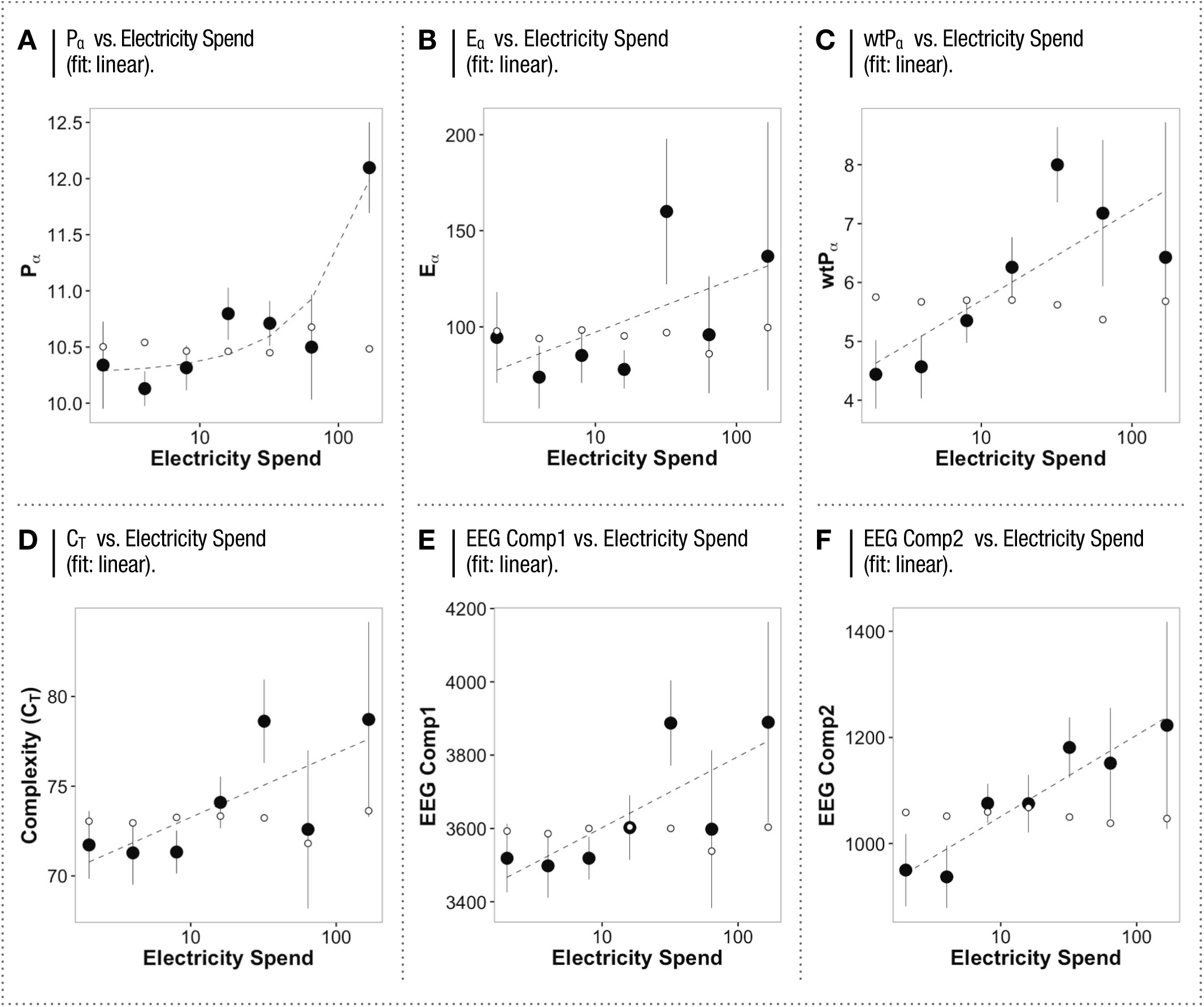
*RELATIONSHIPS OF EEG FEATURES TO ELECTRICITY SPEND*

**Sup. Figure 7.**
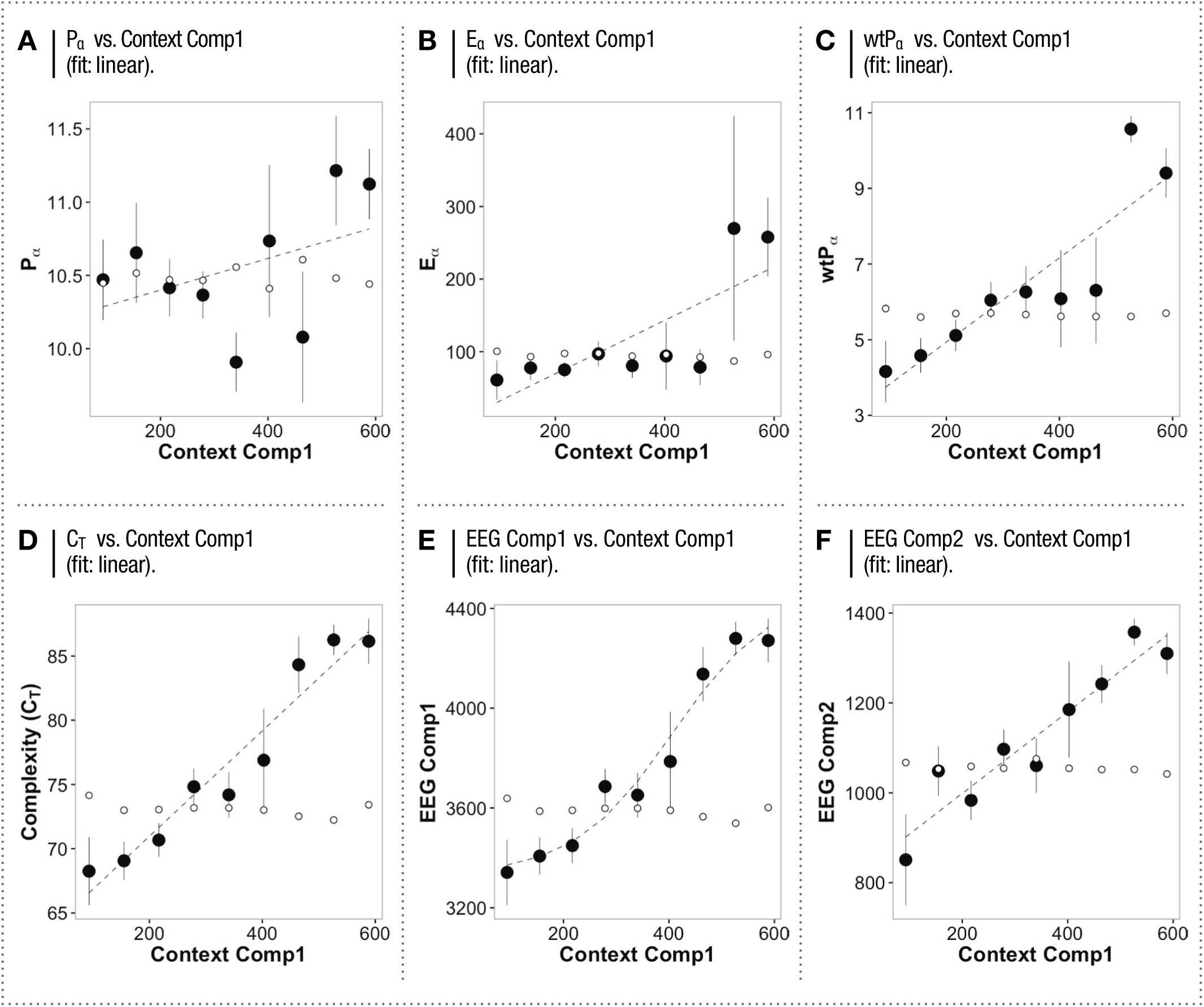
*RELATIONSHIPS OF EEG FEATURES TO CONTEXT COMP1*

**Sup. Figure 8.**
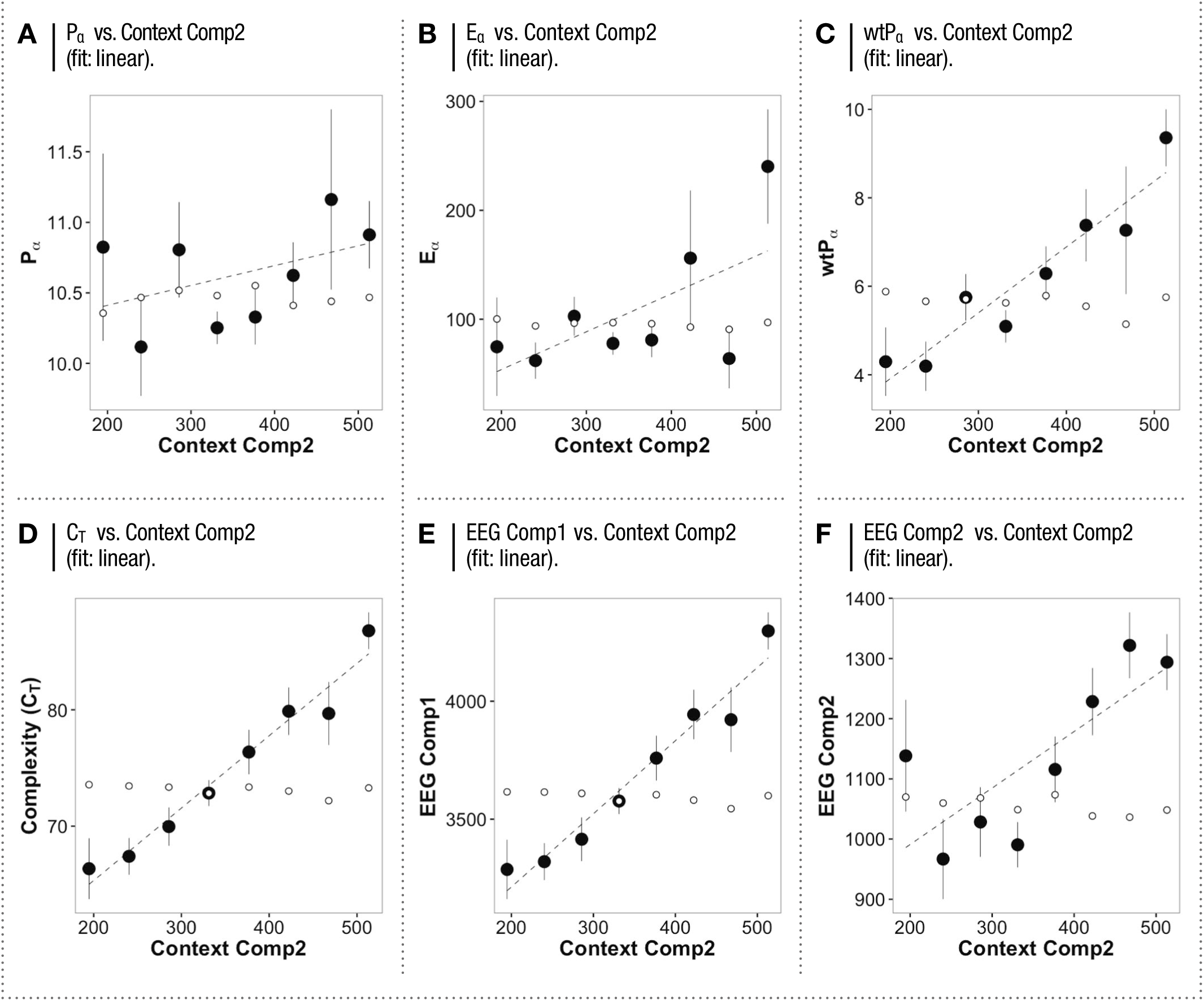
*RELATIONSHIPS OF EEG FEATURES TO CONTEXT COMP2*

**Sup. Table 1.**
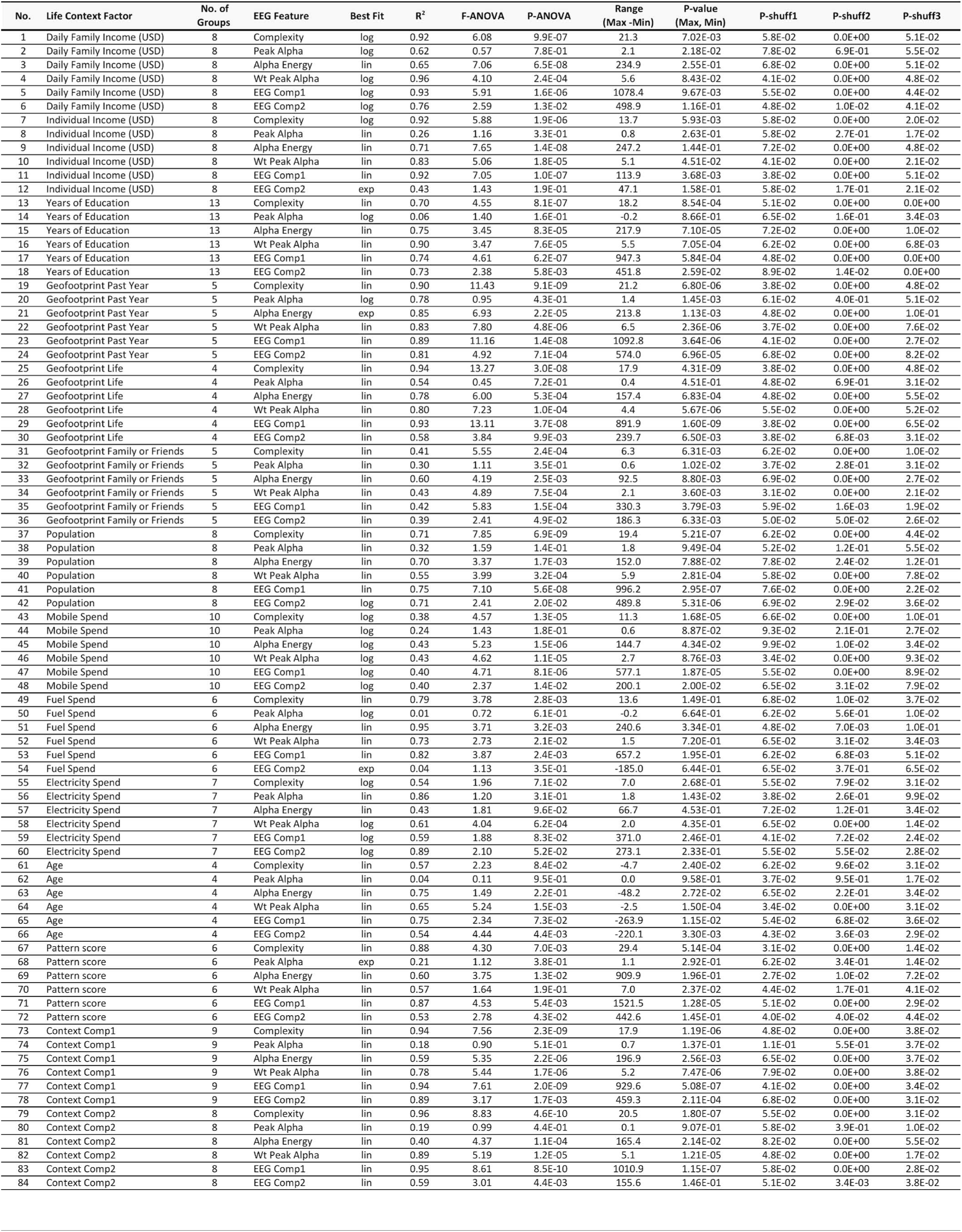

| No. | Life Context Factor | No. of Groups | EEG Feature | Best Fit | R<sup>2</sup> | F-ANOVA | P-ANOVA | Range (Max -Min) | P-value (Max, Min) | P-shuff1 | P-shuff2 | P-shuff3 |
| --- | --- | --- | --- | --- | --- | --- | --- | --- | --- | --- | --- | --- |
| 1 | Daily Family Income (USD) | 8 | Complexity | log | 0.92 | 6.08 | 9.9E-07 | 21.3 | 7.02E-03 | 5.8E-02 | 0.0E+00 | 5.1E-02 |
| 2 | Daily Family Income (USD) | 8 | Peak Alpha | log | 0.62 | 0.57 | 7.8E-01 | 2.1 | 2.18E-02 | 7.8E-02 | 6.9E-01 | 5.5E-02 |
| 3 | Daily Family Income (USD) | 8 | Alpha Energy | lin | 0.65 | 7.06 | 6.5E-08 | 234.9 | 2.55E-01 | 6.8E-02 | 0.0E+00 | 5.1E-02 |
| 4 | Daily Family Income (USD) | 8 | Wt Peak Alpha | log | 0.96 | 4.10 | 2.4E-04 | 5.6 | 8.43E-02 | 4.1E-02 | 0.0E+00 | 4.8E-02 |
| 5 | Daily Family Income (USD) | 8 | EEG Comp1 | log | 0.93 | 5.91 | 1.6E-06 | 1078.4 | 9.67E-03 | 5.5E-02 | 0.0E+00 | 4.4E-02 |
| 6 | Daily Family Income (USD) | 8 | EEG Comp2 | log | 0.76 | 2.59 | 1.3E-02 | 498.9 | 1.16E-01 | 4.8E-02 | 1.0E-02 | 4.1E-02 |
| 7 | Individual Income (USD) | 8 | Complexity | log | 0.92 | 5.88 | 1.9E-06 | 13.7 | 5.93E-03 | 5.8E-02 | 0.0E+00 | 2.0E-02 |
| 8 | Individual Income (USD) | 8 | Peak Alpha | lin | 0.26 | 1.16 | 3.3E-01 | 0.8 | 2.63E-01 | 5.8E-02 | 2.7E-01 | 1.7E-02 |
| 9 | Individual Income (USD) | 8 | Alpha Energy | lin | 0.71 | 7.65 | 1.4E-08 | 247.2 | 1.44E-01 | 7.2E-02 | 0.0E+00 | 4.8E-02 |
| 10 | Individual Income (USD) | 8 | Wt Peak Alpha | lin | 0.83 | 5.06 | 1.8E-05 | 5.1 | 4.51E-02 | 4.1E-02 | 0.0E+00 | 2.1E-02 |
| 11 | Individual Income (USD) | 8 | EEG Comp1 | lin | 0.92 | 7.05 | 1.0E-07 | 113.9 | 3.68E-03 | 3.8E-02 | 0.0E+00 | 5.1E-02 |
| 12 | Individual Income (USD) | 8 | EEG Comp2 | exp | 0.43 | 1.43 | 1.9E-01 | 47.1 | 1.58E-01 | 5.8E-02 | 1.7E-01 | 2.1E-02 |
| 13 | Years of Education | 13 | Complexity | lin | 0.70 | 4.55 | 8.1E-07 | 18.2 | 8.54E-04 | 5.1E-02 | 0.0E+00 | 0.0E+00 |
| 14 | Years of Education | 13 | Peak Alpha | log | 0.06 | 1.40 | 1.6E-01 | -0.2 | 8.66E-01 | 6.5E-02 | 1.6E-01 | 3.4E-03 |
| 15 | Years of Education | 13 | Alpha Energy | lin | 0.75 | 3.45 | 8.3E-05 | 217.9 | 7.10E-05 | 7.2E-02 | 0.0E+00 | 1.0E-02 |
| 16 | Years of Education | 13 | Wt Peak Alpha | lin | 0.90 | 3.47 | 7.6E-05 | 5.5 | 7.05E-04 | 6.2E-02 | 0.0E+00 | 6.8E-03 |
| 17 | Years of Education | 13 | EEG Comp1 | lin | 0.74 | 4.61 | 6.2E-07 | 947.3 | 5.84E-04 | 4.8E-02 | 0.0E+00 | 0.0E+00 |
| 18 | Years of Education | 13 | EEG Comp2 | lin | 0.73 | 2.38 | 5.8E-03 | 451.8 | 2.59E-02 | 8.9E-02 | 1.4E-02 | 0.0E+00 |
| 19 | Geofootprint Past Year | 5 | Complexity | lin | 0.90 | 11.43 | 9.1E-09 | 21.2 | 6.80E-06 | 3.8E-02 | 0.0E+00 | 4.8E-02 |
| 20 | Geofootprint Past Year | 5 | Peak Alpha | log | 0.78 | 0.95 | 4.3E-01 | 1.4 | 1.45E-03 | 6.1E-02 | 4.0E-01 | 5.1E-02 |
| 21 | Geofootprint Past Year | 5 | Alpha Energy | exp | 0.85 | 6.93 | 2.2E-05 | 213.8 | 1.13E-03 | 4.8E-02 | 0.0E+00 | 1.0E-01 |
| 22 | Geofootprint Past Year | 5 | Wt Peak Alpha | lin | 0.83 | 7.80 | 4.8E-06 | 6.5 | 2.36E-06 | 3.7E-02 | 0.0E+00 | 7.6E-02 |
| 23 | Geofootprint Past Year | 5 | EEG Comp1 | lin | 0.89 | 11.16 | 1.4E-08 | 1092.8 | 3.64E-06 | 4.1E-02 | 0.0E+00 | 2.7E-02 |
| 24 | Geofootprint Past Year | 5 | EEG Comp2 | lin | 0.81 | 4.92 | 7.1E-04 | 574.0 | 6.96E-05 | 6.8E-02 | 0.0E+00 | 8.2E-02 |
| 25 | Geofootprint Life | 4 | Complexity | lin | 0.94 | 13.27 | 3.0E-08 | 17.9 | 4.31E-09 | 3.8E-02 | 0.0E+00 | 4.8E-02 |
| 26 | Geofootprint Life | 4 | Peak Alpha | lin | 0.54 | 0.45 | 7.2E-01 | 0.4 | 4.51E-01 | 4.8E-02 | 6.9E-01 | 3.1E-02 |
| 27 | Geofootprint Life | 4 | Alpha Energy | lin | 0.78 | 6.00 | 5.3E-04 | 157.4 | 6.83E-04 | 4.8E-02 | 0.0E+00 | 5.5E-02 |
| 28 | Geofootprint Life | 4 | Wt Peak Alpha | lin | 0.80 | 7.23 | 1.0E-04 | 4.4 | 5.67E-06 | 5.5E-02 | 0.0E+00 | 5.2E-02 |
| 29 | Geofootprint Life | 4 | EEG Comp1 | lin | 0.93 | 13.11 | 3.7E-08 | 891.9 | 1.60E-09 | 3.8E-02 | 0.0E+00 | 6.5E-02 |
| 30 | Geofootprint Life | 4 | EEG Comp2 | lin | 0.58 | 3.84 | 9.9E-03 | 239.7 | 6.50E-03 | 3.8E-02 | 6.8E-03 | 3.1E-02 |
| 31 | Geofootprint Family or Friends | 5 | Complexity | lin | 0.41 | 5.55 | 2.4E-04 | 6.3 | 6.31E-03 | 6.2E-02 | 0.0E+00 | 1.0E-02 |
| 32 | Geofootprint Family or Friends | 5 | Peak Alpha | lin | 0.30 | 1.11 | 3.5E-01 | 0.6 | 1.02E-02 | 3.7E-02 | 2.8E-01 | 3.1E-02 |
| 33 | Geofootprint Family or Friends | 5 | Alpha Energy | lin | 0.60 | 4.19 | 2.5E-03 | 92.5 | 8.80E-03 | 6.9E-02 | 0.0E+00 | 2.7E-02 |
| 34 | Geofootprint Family or Friends | 5 | Wt Peak Alpha | lin | 0.43 | 4.89 | 7.5E-04 | 2.1 | 3.60E-03 | 3.1E-02 | 0.0E+00 | 2.1E-02 |
| 35 | Geofootprint Family or Friends | 5 | EEG Comp1 | lin | 0.42 | 5.83 | 1.5E-04 | 330.3 | 3.79E-03 | 5.9E-02 | 1.6E-03 | 1.9E-02 |
| 36 | Geofootprint Family or Friends | 5 | EEG Comp2 | lin | 0.39 | 2.41 | 4.9E-02 | 186.3 | 6.33E-03 | 5.0E-02 | 5.0E-02 | 2.6E-02 |
| 37 | Population | 8 | Complexity | lin | 0.71 | 7.85 | 6.9E-09 | 19.4 | 5.21E-07 | 6.2E-02 | 0.0E+00 | 4.4E-02 |
| 38 | Population | 8 | Peak Alpha | lin | 0.32 | 1.59 | 1.4E-01 | 1.8 | 9.49E-04 | 5.2E-02 | 1.2E-01 | 5.5E-02 |
| 39 | Population | 8 | Alpha Energy | lin | 0.70 | 3.37 | 1.7E-03 | 152.0 | 7.88E-02 | 7.8E-02 | 2.4E-02 | 1.2E-01 |
| 40 | Population | 8 | Wt Peak Alpha | lin | 0.55 | 3.99 | 3.2E-04 | 5.9 | 2.81E-04 | 5.8E-02 | 0.0E+00 | 7.8E-02 |
| 41 | Population | 8 | EEG Comp1 | lin | 0.75 | 7.10 | 5.6E-08 | 996.2 | 2.95E-07 | 7.6E-02 | 0.0E+00 | 2.2E-02 |
| 42 | Population | 8 | EEG Comp2 | log | 0.71 | 2.41 | 2.0E-02 | 489.8 | 5.31E-06 | 6.9E-02 | 2.9E-02 | 3.6E-02 |
| 43 | Mobile Spend | 10 | Complexity | log | 0.38 | 4.57 | 1.3E-05 | 11.3 | 1.68E-05 | 6.6E-02 | 0.0E+00 | 1.0E-01 |
| 44 | Mobile Spend | 10 | Peak Alpha | log | 0.24 | 1.43 | 1.8E-01 | 0.6 | 8.87E-02 | 9.3E-02 | 2.1E-01 | 2.7E-02 |
| 45 | Mobile Spend | 10 | Alpha Energy | log | 0.43 | 5.23 | 1.5E-06 | 144.7 | 4.34E-02 | 9.9E-02 | 1.0E-02 | 3.4E-02 |
| 46 | Mobile Spend | 10 | Wt Peak Alpha | log | 0.43 | 4.62 | 1.1E-05 | 2.7 | 8.76E-03 | 3.4E-02 | 0.0E+00 | 9.3E-02 |
| 47 | Mobile Spend | 10 | EEG Comp1 | log | 0.40 | 4.71 | 8.1E-06 | 577.1 | 1.87E-05 | 5.5E-02 | 0.0E+00 | 8.9E-02 |
| 48 | Mobile Spend | 10 | EEG Comp2 | log | 0.40 | 2.37 | 1.4E-02 | 200.1 | 2.00E-02 | 6.5E-02 | 3.1E-02 | 7.9E-02 |
| 49 | Fuel Spend | 6 | Complexity | lin | 0.79 | 3.78 | 2.8E-03 | 13.6 | 1.49E-01 | 6.8E-02 | 1.0E-02 | 3.7E-02 |
| 50 | Fuel Spend | 6 | Peak Alpha | log | 0.01 | 0.72 | 6.1E-01 | -0.2 | 6.64E-01 | 6.2E-02 | 5.6E-01 | 1.0E-02 |
| 51 | Fuel Spend | 6 | Alpha Energy | lin | 0.95 | 3.71 | 3.2E-03 | 240.6 | 3.34E-01 | 4.8E-02 | 7.0E-03 | 1.0E-01 |
| 52 | Fuel Spend | 6 | Wt Peak Alpha | lin | 0.73 | 2.73 | 2.1E-02 | 1.5 | 7.20E-01 | 6.5E-02 | 3.1E-02 | 3.4E-03 |
| 53 | Fuel Spend | 6 | EEG Comp1 | lin | 0.82 | 3.87 | 2.4E-03 | 657.2 | 1.95E-01 | 6.2E-02 | 6.8E-03 | 5.1E-02 |
| 54 | Fuel Spend | 6 | EEG Comp2 | exp | 0.04 | 1.13 | 3.5E-01 | -185.0 | 6.44E-01 | 6.5E-02 | 3.7E-01 | 6.5E-02 |
| 55 | Electricity Spend | 7 | Complexity | log | 0.54 | 1.96 | 7.1E-02 | 7.0 | 2.68E-01 | 5.5E-02 | 7.9E-02 | 3.1E-02 |
| 56 | Electricity Spend | 7 | Peak Alpha | lin | 0.86 | 1.20 | 3.1E-01 | 1.8 | 1.43E-02 | 3.8E-02 | 2.6E-01 | 9.9E-02 |
| 57 | Electricity Spend | 7 | Alpha Energy | lin | 0.43 | 1.81 | 9.6E-02 | 66.7 | 4.53E-01 | 7.2E-02 | 1.2E-01 | 3.4E-02 |
| 58 | Electricity Spend | 7 | Wt Peak Alpha | log | 0.61 | 4.04 | 6.2E-04 | 2.0 | 4.35E-01 | 6.5E-02 | 0.0E+00 | 1.4E-02 |
| 59 | Electricity Spend | 7 | EEG Comp1 | log | 0.59 | 1.88 | 8.3E-02 | 371.0 | 2.46E-01 | 4.1E-02 | 7.2E-02 | 2.4E-02 |
| 60 | Electricity Spend | 7 | EEG Comp2 | log | 0.89 | 2.10 | 5.2E-02 | 273.1 | 2.33E-01 | 5.5E-02 | 5.5E-02 | 2.8E-02 |
| 61 | Age | 4 | Complexity | lin | 0.57 | 2.23 | 8.4E-02 | -4.7 | 2.40E-02 | 6.2E-02 | 9.6E-02 | 3.1E-02 |
| 62 | Age | 4 | Peak Alpha | lin | 0.04 | 0.11 | 9.5E-01 | 0.0 | 9.58E-01 | 3.7E-02 | 9.5E-01 | 1.7E-02 |
| 63 | Age | 4 | Alpha Energy | lin | 0.75 | 1.49 | 2.2E-01 | -48.2 | 2.72E-02 | 6.5E-02 | 2.2E-01 | 3.4E-02 |
| 64 | Age | 4 | Wt Peak Alpha | lin | 0.65 | 5.24 | 1.5E-03 | -2.5 | 1.50E-04 | 3.4E-02 | 0.0E+00 | 3.1E-02 |
| 65 | Age | 4 | EEG Comp1 | lin | 0.75 | 2.34 | 7.3E-02 | -263.9 | 1.15E-02 | 5.4E-02 | 6.8E-02 | 3.6E-02 |
| 66 | Age | 4 | EEG Comp2 | lin | 0.54 | 4.44 | 4.4E-03 | -220.1 | 3.30E-03 | 4.3E-02 | 3.6E-03 | 2.9E-02 |
| 67 | Pattern score | 6 | Complexity | lin | 0.88 | 4.30 | 7.0E-03 | 29.4 | 5.14E-04 | 3.1E-02 | 0.0E+00 | 1.4E-02 |
| 68 | Pattern score | 6 | Peak Alpha | exp | 0.21 | 1.12 | 3.8E-01 | 1.1 | 2.92E-01 | 6.2E-02 | 3.4E-01 | 1.4E-02 |
| 69 | Pattern score | 6 | Alpha Energy | lin | 0.60 | 3.75 | 1.3E-02 | 909.9 | 1.96E-01 | 2.7E-02 | 1.0E-02 | 7.2E-02 |
| 70 | Pattern score | 6 | Wt Peak Alpha | lin | 0.57 | 1.64 | 1.9E-01 | 7.0 | 2.37E-02 | 4.4E-02 | 1.7E-01 | 4.1E-02 |
| 71 | Pattern score | 6 | EEG Comp1 | lin | 0.87 | 4.53 | 5.4E-03 | 1521.5 | 1.28E-05 | 5.1E-02 | 0.0E+00 | 2.9E-02 |
| 72 | Pattern score | 6 | EEG Comp2 | lin | 0.53 | 2.78 | 4.3E-02 | 442.6 | 1.45E-01 | 4.0E-02 | 4.0E-02 | 4.4E-02 |
| 73 | Context Comp1 | 9 | Complexity | lin | 0.94 | 7.56 | 2.3E-09 | 17.9 | 1.19E-06 | 4.8E-02 | 0.0E+00 | 3.8E-02 |
| 74 | Context Comp1 | 9 | Peak Alpha | lin | 0.18 | 0.90 | 5.1E-01 | 0.7 | 1.37E-01 | 1.1E-01 | 5.5E-01 | 3.7E-02 |
| 75 | Context Comp1 | 9 | Alpha Energy | lin | 0.59 | 5.35 | 2.2E-06 | 196.9 | 2.56E-03 | 6.5E-02 | 0.0E+00 | 3.7E-02 |
| 76 | Context Comp1 | 9 | Wt Peak Alpha | lin | 0.78 | 5.44 | 1.7E-06 | 5.2 | 7.47E-06 | 7.9E-02 | 0.0E+00 | 3.8E-02 |
| 77 | Context Comp1 | 9 | EEG Comp1 | lin | 0.94 | 7.61 | 2.0E-09 | 929.6 | 5.08E-07 | 4.1E-02 | 0.0E+00 | 3.4E-02 |
| 78 | Context Comp1 | 9 | EEG Comp2 | lin | 0.89 | 3.17 | 1.7E-03 | 459.3 | 2.11E-04 | 6.8E-02 | 0.0E+00 | 3.1E-02 |
| 79 | Context Comp2 | 8 | Complexity | lin | 0.96 | 8.83 | 4.6E-10 | 20.5 | 1.80E-07 | 5.5E-02 | 0.0E+00 | 3.1E-02 |
| 80 | Context Comp2 | 8 | Peak Alpha | lin | 0.19 | 0.99 | 4.4E-01 | 0.1 | 9.07E-01 | 5.8E-02 | 3.9E-01 | 1.0E-02 |
| 81 | Context Comp2 | 8 | Alpha Energy | lin | 0.40 | 4.37 | 1.1E-04 | 165.4 | 2.14E-02 | 8.2E-02 | 0.0E+00 | 5.5E-02 |
| 82 | Context Comp2 | 8 | Wt Peak Alpha | lin | 0.89 | 5.19 | 1.2E-05 | 5.1 | 1.21E-05 | 4.8E-02 | 0.0E+00 | 1.7E-02 |
| 83 | Context Comp2 | 8 | EEG Comp1 | lin | 0.95 | 8.61 | 8.5E-10 | 1010.9 | 1.15E-07 | 5.8E-02 | 0.0E+00 | 2.8E-02 |
| 84 | Context Comp2 | 8 | EEG Comp2 | lin | 0.59 | 3.01 | 4.4E-03 | 155.6 | 1.46E-01 | 5.1E-02 | 3.4E-03 | 3.8E-02 |

